# Dual fragmentation via collision-induced and oxygen attachment dissociations using water and its radicals for C=C position-resolved lipidomics

**DOI:** 10.1101/2024.10.31.621229

**Authors:** Hiroaki Takeda, Mami Okamoto, Hidenori Takahashi, Bujinlkham Buyantogtokh, Noriyuki Kishi, Hideyuki Okano, Hiroyuki Kamiguchi, Hiroshi Tsugawa

**Affiliations:** Department of Biotechnology and Life Science, Tokyo University of Agriculture and Technology, 2-24-16 Naka-cho, Koganei, Tokyo 184-8588, Japan; RIKEN Center for Brain Science, 2-1 Hirosawa, Wako, Saitama 351-0106, Japan; Shimadzu Corporation, 1 Nishinokyo-Kuwabaracho Nakagyo-ku, Kyoto, 604-8511, Japan; Department of Physiology, Keio University School of Medicine, 35 Shinanomachi, Shinjuku-ku, Tokyo 160-8582, Japan; RIKEN Center for Sustainable Resource Science, 1-7-22 Suehiro-cho, Tsurumi-ku, Yokohama, Kanagawa 230-0045, Japan; RIKEN Center for Integrative Medical Sciences, 1-7-22 Suehiro-cho, Tsurumi-ku, Yokohama, Kanagawa 230-0045, Japan; Graduate School of Medical Life Science, Yokohama City University, 1-7-29 Suehiro-cho, Tsurumi-ku, Yokohama, Kanagawa 230-0045, Japan

**Author notes:** **Corresponding Authors**: Hiroaki Takeda (H. Takeda), Hiroshi Tsugawa (H. Tsugawa).

## Abstract

Oxygen attachment dissociation (OAD) is a tandem mass spectrometry (MS/MS) technique used to annotate the positions of double bonds (C=C) in complex lipids. Although OAD has been used for untargeted lipidomics, its availability has been limited to the positive-ion mode, requiring the independent use of a collision-induced dissociation (CID) method. In this study, we demonstrated the OAD-MS/MS technique in the negative-ion mode for profiling phosphatidylserines, phosphatidylglycerols, phosphatidylinositols, and sulfatides, where the fragmentation mechanism remained consistent with that in the positive-ion mode. Furthermore, we proposed optimal conditions for the simultaneous acquisition of CID– and OAD-specific fragment ions, termed OAciD. In the collision cell for OAD, oxygen atoms and hydroxy radicals facilitate C=C position-specific fragmentation, while residual water vapor induces cleavage of low-energy covalent bonds, such as ester and peptide bonds, at higher collision energy values, preserving OAD-specific ions under high collision energy conditions. Finally, theoretical fragment ions were implemented in MS-DIAL 5 to accelerate C=C position-resolved untargeted lipidomics. The OAciD methodology was applied to lipid profiling of five marmoset brain regions: the frontal lobe, hippocampus, midbrain, cerebellum, and medulla. Region-specific marmoset lipidomes were characterized with C=C positional information, where the ratios of C=C positional isomers such as delta 9– and delta 11 of fatty acid 18:1 in phosphatidylcholine were also estimated using OAciD-MS/MS. In addition, we characterized the profiles of polyunsaturated fatty acid-containing complex lipids with C=C positional information, where lipids containing omega-3 fatty acids were enriched in the cerebellum, while those containing omega-6 fatty acids were more abundant in the hippocampus and frontal lobe.

## Introduction

Lipids are essential components of cell membranes, signal transduction pathways, and energy storage materials.^1–3^ They consist of a backbone (e.g., glycerol, sphingosine, or sterol), a polar head group (e.g., phosphocholine), and fatty acyl chains that vary in length and number of unsaturated bonds. These fatty acyl chains exist in various isomers, differing in double bond positions, *sn*-positions, and cis-trans configurations, contributing to the approximately 50,000 lipids registered in the LIPID MAPS Structure Database.^4^ Disruption in the quality and quantity of lipid diversity is linked to cellular homeostasis dysfunction, leading to disease in animals.^5^

Recently, untargeted lipidomics has gained attention for capturing lipid alterations and understanding the link between lipid metabolism and phenotypes.^6^ Electrospray ionization (ESI) coupled with collision-induced dissociation (CID)-based tandem mass spectrometry (MS/MS) is commonly used to characterize lipid structures. While CID-based fragmentation enables the molecular species-level annotation of lipids, such as determining polar head groups (e.g., phosphocholine of phosphatidylcholine (PC) in the positive ion mode) and total fatty acyl compositions (e.g., fatty acyl chains in the negative ion mode), it does not provide information on C=C positions in fatty acyl chains.

To address this, recent analytical methods have been developed to elucidate the detailed structure of fatty acyl chains in complex lipids. Charge-switch derivatization with *N*-(4-aminomethylphenyl) pyridinium not only enhances the sensitivity of fatty acids (FAs) but also provides multiple diagnostic fragment ions, enabling the annotation of C=C positions.^7^ Additionally, the Paternò-Büchi reaction and epoxidation using *meta*-chloroperoxybenzoic acid are effective pretreatment methods for determining double bond positions.^8,9^ Furthermore, fragmentation techniques such as ultraviolet photodissociation,^10^ ozone-induced dissociation (OzID),^11,12^ oxygen attachment dissociation (OAD),^13,14^ and electron-activated dissociation (EAD)^15–17^ have emerged, allowing for in-depth structural analysis without requiring derivatization or specialized equipment. Among these, EAD is a commercialized method available in the positive ion mode, providing insights into double bonds, *sn*-positions, and potentially *E/Z* isomers.^15–17^ This method allows for detailed lipid structure elucidation for abundant lipids at concentrations of 1 µg/mL or more. Moreover, the EAD technique requires metal ions, such as sodium ions, to facilitate the structural elucidation of negatively charged glycerophospholipids like phosphatidylinositol (PI), phosphatidylglycerol (PG), and phosphatidylserine (PS), using non-volatile sodium salts.

OAD-MS/MS, now commercially available, employs radical-induced fragmentation using atomic oxygen (O) and/or hydroxyl radicals (OH·) generated by the microwave irradiation of water vapor (H_2_O). These reactive species interact with the electrons in double bonds (R1-C=C-R2), cleaving C=C bonds between R1 and C, and C and R2. OAD’s fragmentation patterns are similar to those of OzID, acquiring C=C position-specific fragment ions, but OAD is easier to handle and safer than ozone, a strong oxidant. In our previous study, we demonstrated C=C position-resolved lipidomics in positive ion mode by integrating molecular species-level annotation obtained from ESI(+)– and ESI(−)-CID-MS/MS techniques.^18^ For example, “PC 16:0_18:1” was first annotated by CID-MS/MS, and the C=C position was determined using OAD-MS/MS, resulting in the final annotation “PC 16:0_18:1(Δ9)”. While the successful annotation of cationic lipids, such as PC and phosphatidylethanolamine (PE), was shown, the OAD technique can be applied in both positive and negative ion modes. CID-specific fragment ions related to polar head groups and FA composition can also be obtained under OAD’s collision cell conditions.

In this study, we investigated OAD mass fragmentation in both positive and negative ion modes to reveal C=C structural isomers, including negatively charged lipid molecules. We also explored a collision cell condition that generated both CID– and OAD-specific fragment ions using H_2_O and its radicals, termed OAciD, which enabled in-depth structural elucidation of lipids with a single analytical method. Finally, we developed and implemented an algorithm for automatic lipid annotation based on the quality of the acquired product ion spectra in MS-DIAL 5 software.^19,20^ The accuracy of the automated annotation was evaluated using an UltimateSPLASH mixture containing 69 synthetic lipid standards. Using this advanced technique, we performed untargeted lipidomics on the marmoset brain, characterizing the unique lipidomes of the frontal lobe, hippocampus, midbrain, cerebellum, and medulla.

## Results

### Extension of lipid coverage by using both positive and negative ion modes

To extend lipid coverage, we evaluated the fragmentation patterns and efficiency of OAD-MS/MS in the negative ion mode, a domain that has not been thoroughly explored for lipids. Additionally, we examined sodium adduct ions, as recommended in a previous lipidomics study under EAD conditions,^17^ to efficiently determine the C=C positions of negatively charged lipids such as PG and PI. The use of organic solvents containing sodium acetate, introduced post-column, increases the formation and sensitivity of sodium adduct ions.^21^ However, employing sodium salts presents several challenges, including increased background noise in the total ion chromatogram, reduced sensitivity and reproducibility of conventional proton and ammonium adduct ions, and mass spectrometer contamination due to sodium’s nonvolatility. We compared the sensitivity of the MS1 peaks under three ionization conditions: conventional proton or ammonium adduct ions in the positive ion mode ([M+H]^+^ or [M+NH_4_]^+^), deprotonated or acetate adduct ions in the negative ion mode ([M−H]^−^ or [M+CH_3_COO]^−^), and sodium adduct ions in the positive ion mode ([M+Na]^+^). The column eluent (600 μL/min total) was mixed with 50 μL/min of 50% methanol containing 200 μM sodium acetate post-column to form the sodium adduct ion (Figure S1a). Although the addition of sodium acetate solution increased background noise (Figure S1b), the peak area reproducibility of proton, ammonium, and sodium adduct ions was within 10% for each lipid subclass when 50 μL/min of sodium solution was mixed post-column (Figure S1c). Among lipid subclasses, diacylglycerol (DG) and ceramide (Cer) showed improved sensitivities with the sodium adduct form (Figure S1d). In contrast, the negative ion mode provided higher sensitivity for detecting PG and PI (Figure S1d).

We further evaluated the sensitivity of the MS/MS peaks using a product ion scan in which appropriate precursor ions were selected based on the full MS scan data (Table S1). In the fragmentation of FA 18:1(Δ9) fatty acyl moieties, neutral losses of 96.13 Da and 138.14 Da, corresponding to the cleavages of Δ10-Δ11 and Δ8-Δ9 positions, respectively, were observed (Figure 1a).^18^ No differences in fragmentation patterns were detected when comparing MS/MS spectra between positive and negative ion modes, indicating that the fragmentation mechanism was unaffected by polarity differences (Figure 1b). However, the ratio of fragmentation efficiency between positive and negative ion modes varied across lipid subclasses. The highest MS/MS peaks for lyso-PC (LPC), lyso-PE (LPE), PC, sphingomyelin (SM), DG, and triacylglycerol (TG) were observed in sodium adduct forms, whereas better sensitivities were achieved in the negative ion mode for PG, PI, and Cer (Figure S1d). The high fragmentation efficiency of LPC, LPE, PC, SM, DG, and TG in the sodium adduct form is likely due to the strong charge bias introduced by metal ions like sodium, compared to proton or ammonium ions.

**Figure 1.**
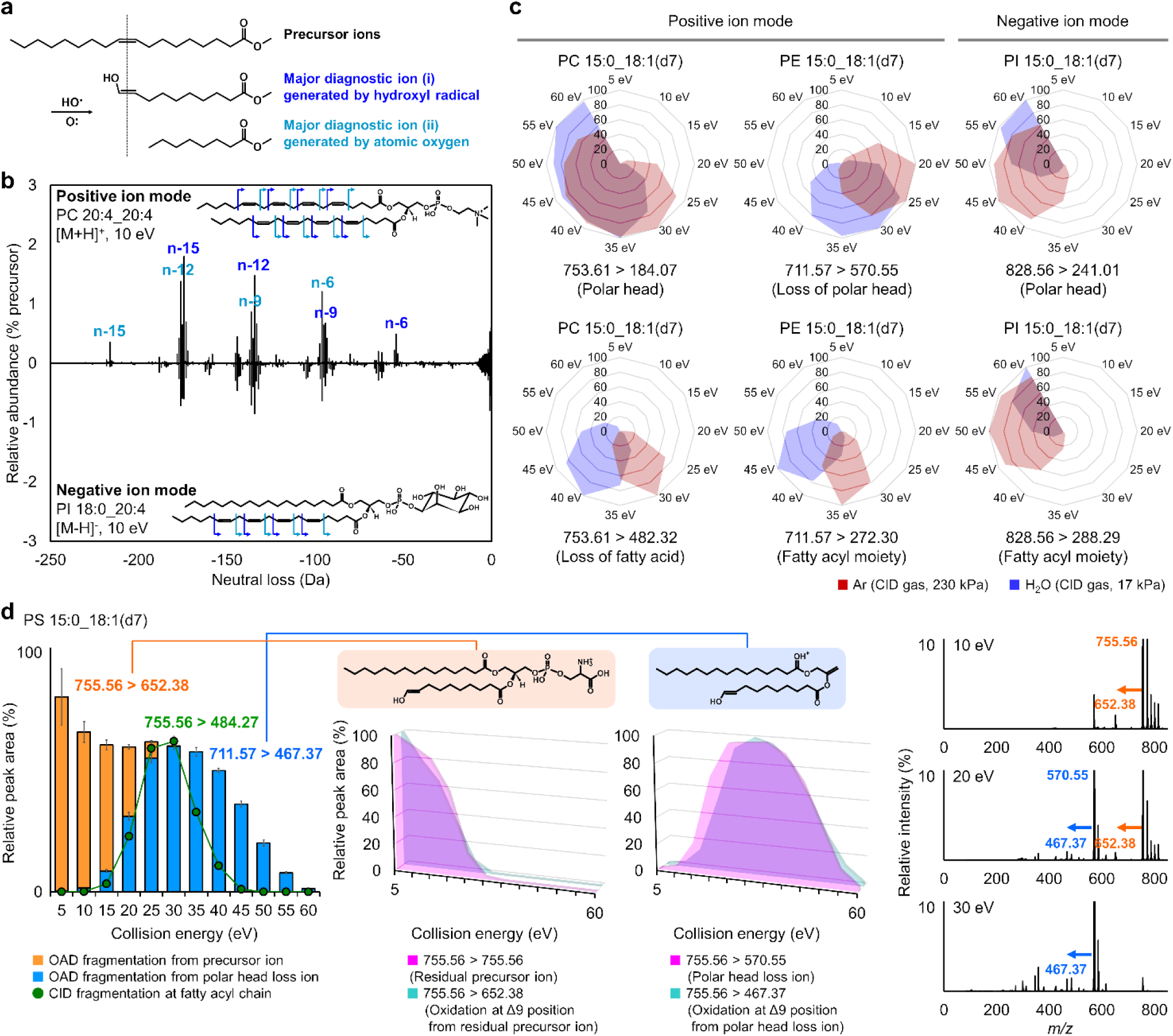
Characteristics of OAciD-MS/MS. (**a**) Major fragmentation of OAD-MS/MS. (**b**) Fragmentation difference between positive and negative ion modes. (**c**) Appropriate collision energy in Ar gas and H_2_O vapor. The area value of MS2 peak was normalized to the maximum peak detected under the collision energy range of 5–60 eV. All lipid classes are summarized in Figure S2. (**d**) Fragmentation at the C=C position by increasing the collision energy. In the left panel, the orange and blue bars represent the MS2 peak areas of fragments at the C=C position, derived from the residual precursor and polar head loss ions, respectively. The green line plot indicates fragments from fatty acyl chains, also observed in CID mode using Ar gas. Error bars represent the standard deviations of four analytical replicates. A summary of all lipid classes is provided in Figure S3. The middle panels show the MS2 peak areas of fragments at the C=C position from residual precursor and polar head loss ions. A summary of all lipid classes is provided in Figure S4. The right panels display the MS/MS spectra at 10, 20, and 30 eV.

Nevertheless, we chose to use protons and ammonium ions, because data-dependent MS/MS acquisition in untargeted lipidomics is more efficient when the signal-to-noise ratio is higher in full scan MS1. Based on our results, an ammonium acetate-containing LC solvent, which efficiently detects protons or ammonium ions, facilitates data-dependent MS/MS-based lipidomics for screening purposes. Meanwhile, using sodium adduct forms in targeted MS/MS mode would accelerate in-depth structural elucidation. In this study, as our aim was to develop a lipidomics pipeline using data-dependent OAciD-MS/MS, we optimized the analytical method using proton and ammonium ions, enabling lipid quantification through MS1 peaks and annotation via data-dependent MS/MS.

### Dual fragmentation of CID and OAD using H_2_O and its radicals

Fragment ions of polar head groups and fatty acyl moieties are essential for annotating lipid molecules using MS/MS. For instance, the phosphorylcholine fragment ion (*m/z* 184.07) detected in the positive ion mode is useful for annotating LPC, PC, and SM. However, the MS/MS intensity of these polar head-specific fragment ions was insufficient in previous OAD-MS/MS systems because the collision cell utilized only 5–10 eV, which is adequate for ion transfer but not for efficient fragmentation. We hypothesized that by increasing the collision energy within the collision cell, fragment ions from simultaneous OAD and CID reactions could be observed, as typical lipid product ion spectra contain a portion of unreacted ions (i.e., precursor ions).

Inert gases, such as argon (Ar) and nitrogen, are typically used in CID-MS/MS to minimize unwanted reactions in collision cells. However, OAD-MS/MS employs O and/or OH radicals generated by microwave irradiation of H_2_O vapor for radical reactions. Therefore, it is necessary to obtain fragment ions of polar head groups and fatty acyl moieties using residual H_2_O vapor instead of inert gases in this system. We first investigated the differences in fragmentation patterns between H_2_O vapor and Ar. The results showed that the appropriate collision energy for generating product ions related to polar head groups and FA chains was on average 10 eV higher with H_2_O vapor than with Ar gas (Figure 1c and S2), although the fragmentation patterns remained similar under both conditions.

We further explored the sensitivity of fragment ions derived from the C=C position (i.e., OAD reactions) by varying the collision energy from 10 to 60 eV. The product ion of the C=C position-related fragment of PC decreased at collision energies above 20 eV, while both fatty acyl-related fragment ions (*m/z* 482.32) and C=C position-related fragment ions (*m/z* 650.44) were observed at 30–35 eV (Figure S3a). Notably, when comparing the MS2 peak areas of the residual precursor and the C=C position-related fragment ion, the sensitivities were highly correlated with collision energy conditions (Figure S4a), indicating that the sensitivity of C=C position-related fragment ions depends on the ion abundance of the residual precursor. Neutral-loss ions of polar head groups, such as those observed in PE, PG, and PS, were detected with increased collision energy (Figure S2). Two types of C=C position-related fragment ions were observed in these subclasses: one derived from the precursor ion and the other from the product ion generated by the neutral loss of the polar head groups (Figures 1d and S3a). The total peak areas of these oxidative fragments from the residual precursor and neutral loss ions were nearly identical (Figure 1d and S3a). Since the OAD reaction is slower than the CID reaction, OAciD-MS/MS with higher collision energy produces spectra that can be obtained in MS3, enabling sequential fragmentation from the polar-head-related neutral loss ion to the C=C position-specific ion. The strong correlation between the neutral-loss fragments of polar head groups and the oxidative fragments at the C=C position from these neutral-loss fragments further confirms that the OAD reaction does not depend on collision energy (Figure 1d and S4).

Additionally, we found that a collision energy of 30 eV was optimal for generating both C=C position (OAD) and fatty acyl composition (CID)-related fragment ions (Figures 1d and S2–S4). Thus, we used 30 eV as the collision energy for dual fragmentation of OAD and CID (OAciD) in positive ion mode. The collision energy in negative ion mode should be optimized to efficiently detect the PI and PG molecules, as their detection was less effective in the positive ion mode. While higher collision energy is required when annotating the fatty acyl composition of PI (Figure 1c), we also set the collision energy at 30 eV in the negative ion mode analysis, balancing the sensitivity for C=C position (OAD) and fatty acyl composition (CID) in PI molecules (*m/z* 241.01, 0.61 ± 0.13%; *m/z* 288.29, 0.15 ± 0.03%; *m/z* 683.37, 0.18 ± 0.05%; and *m/z* 725.39, 0.37 ± 0.08% from the residual precursor intensity) (Figures S2b and S3b). The MS/MS spectra of PC and PI obtained using this optimized method are shown in Figure 2a. Despite the complexity introduced by H_2_O vapor adduct ions, our findings indicate that using 30 eV in the collision cell enables the simultaneous detection of polar head, FAs, and C=C position-specific fragment ions in a single analysis.

**Figure 2.**
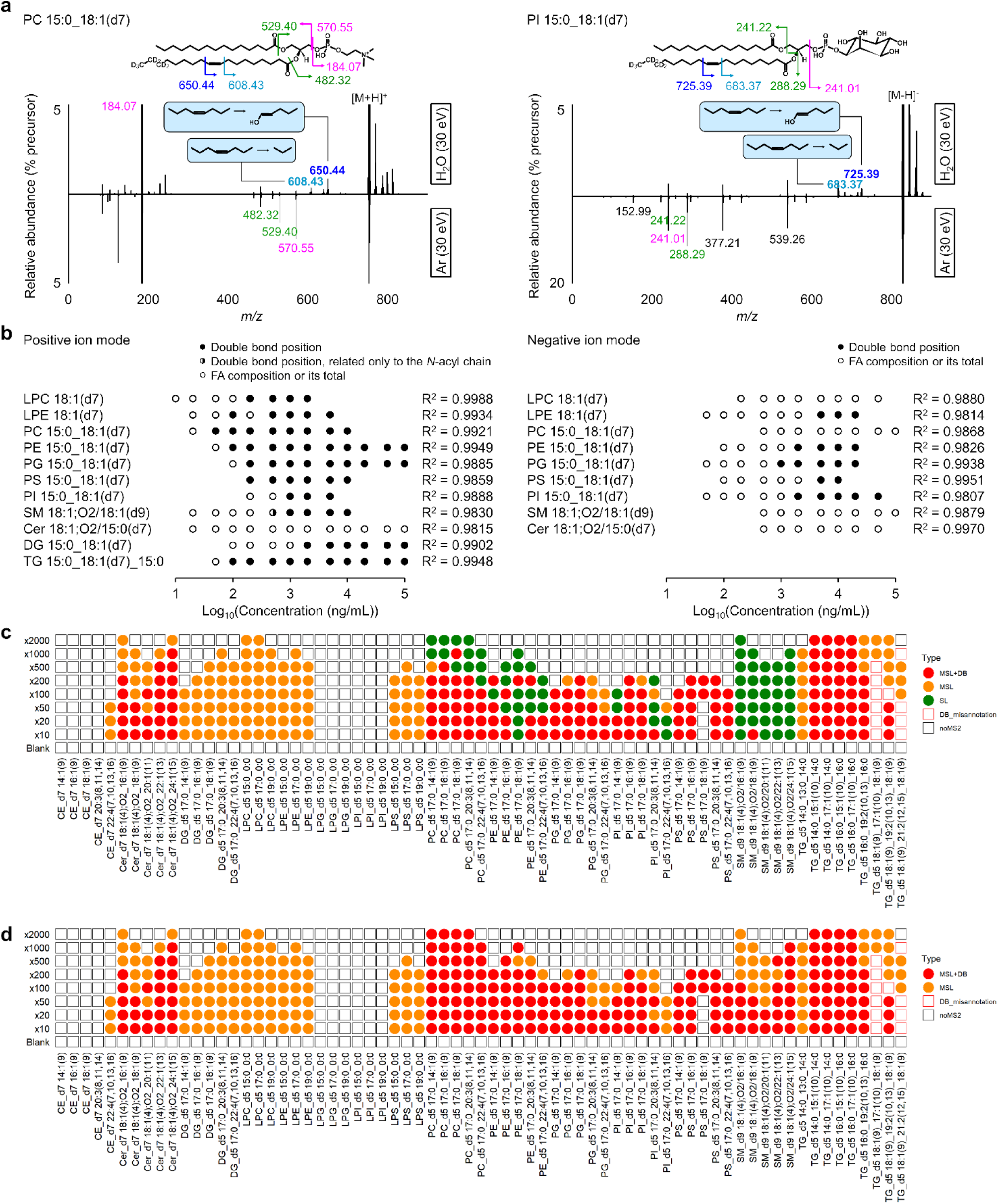
Automated annotation of lipid standards using MS-DIAL 5 software. (**a**) Summary of MS/MS spectra by setting 30 eV of collision energy in both positive and negative ion modes. (**b**) Calibration curve of each lipid subclass in positive and negative ion modes. Data-dependent MS/MS mode was used for evaluation. Circles represent the linear range of the MS1 peak with R^2^ > 0.9800 (Table S1). The closed circles represent concentrations where the C=C position could be determined, with two major fragments detected simultaneously in at least two out of four analyses. The half-filled circle for SM indicates that the C=C position was determined only on the *N*-acyl chain, not the long chain base. Annotation depth was assessed using the EquiSPLASH mixture containing deuterium-labeled standards. (**c**) Automated annotation results of 69 synthetic lipid standards using the optimized MS-DIAL 5 software. The UltimateSPLASH standard mixture was diluted 10, 20, 50, 100, 200, 500, 1000, and 2000 times with MeOH. For example, “x10” indicates a dilution 10 times less concentrated than the original, denoted as “x1”. Colors represent the structural depth of the automatic annotation: red indicates both fatty acyl composition and double bond position (MSL+DB); orange indicates fatty acyl composition only (MSL); green indicates species level annotation only (SL). If the MS/MS spectrum was not assigned to the precursor ion by DDA, a square shape with black color is used (noMS2). If the annotation for double bond characterization was incorrect, a square shape with red color is used (DB_misannotation). The representative annotation was determined as follows: if the same lipid name was annotated in at least two of the four replicates, that name was used as the representative annotation. If the annotation results differed across all four replicates, the lipid with the highest score was adopted as the representative. Lysophospholipids such as LPC, LPE, LPG, lyso-PI (LPI), and lyso-PS (LPS) are out of range for C=C position determination due to their saturated fatty acyl moieties. (**d**) Annotation results using the MS-DIAL 5 program when the MSL annotations were carried out before the C=C positional annotations are performed. The definitions of color and symbol are the same as used in Figure 2c.

### Evaluation of OAciD-MS/MS spectral annotations using MS-DIAL 5

The dynamic range of lipid quantification was investigated using the EquiSPLASH standard containing 100 μg/mL of 13 deuterium-labeled synthetic lipids with a double bond at the Δ9 position except for Cer 18:1;O2/15:0 (Table S2). Owing to the narrow linear range of the fragment ions (less than two digits), the lipids were quantified using MS1 peaks, and their detailed structures were characterized using fragments obtained by data-dependent MS/MS. To increase the number of MS1 peaks that can characterize the C=C position while maintaining the linear range of the MS1 peaks, MS1 and MS2 ion accumulations were turned off and on, respectively, in the Shimadzu MS setting. The double bond position could also be characterized at low concentrations in the linear range using the positive ion mode (Figure 2b). However, the sensitivity of C=C position-related fragment ions in the OAciD-MS/MS spectra in negative ion mode is still challenging, although OAciD-MS/MS works for the PI and PG targeted in this study (Figure 2b). In addition, the MS-DIAL 5 software program was updated for fragment annotation of the OAciD-MS/MS spectra. The program uses a rule-based (decision-tree-based) algorithm, in which the existence of diagnostic ions related to the polar head, fatty acyls, and C=C positions are confirmed. Importantly, MS-DIAL requires molecular species-level (MSL) annotations such as PC 16:0_18:1 to determine the C=C position. Furthermore, the program excluded candidates if the major diagnostic fragment ions related to the C=C position were not detected in the MS/MS spectrum. For example, neutral losses of 96.13 Da (Δ10-Δ11 cleavage) and 138.14 Da (Δ8-Δ9 cleavage) were used for the diagnostic fragments of PC 16:0_18:1(Δ9). Finally, candidates with all diagnostic fragment ions in the MS/MS spectrum were ranked based on the correlation coefficient between the experimental and computationally generated *in silico* MS/MS spectra.

We evaluated the annotation accuracy of MS-DIAL 5 using liquid chromatography-mass spectrometry (LC-MS) data obtained from a dilution series of UltimateSPLASH containing 69 deuterium-labeled synthetic lipids at various concentrations. This standard mixture contained varying numbers of double bonds in the acyl chains of FAs, including polyunsaturated FA (PUFA). Among the 10 lipid subclasses, the fatty acyl compositions and their double bond positions were accurately determined in positive ion mode, except for cholesteryl ester (CE), DG, SM, and some PE and TG molecules (Figure 2c and Figure S5a). Of these, the annotation results of SM, PE, and TG molecules became species level (SL) because the *O*-acyl and *N*-acyl fragment ions used for MSL annotations were difficult to detect in the MS/MS spectra, whereas the C=C-specific fragment ions were clearly detected. The sensitivity of the fragment ion (*m/z* 264.27) for *N*-acyl chain determination of the SM lipid subclass was particularly low at 30 eV (Figure S2a). Thus, we further evaluated the MS-DIAL 5 program in cases where MSL annotations for all lipid subclasses were accomplished before C=C position was determined (Figure 2d). The results indicated that MS-DIAL 5 correctly annotated the C=C positions of phospholipids and SM. The results of misannotations in several TG molecules indicated that the interpretation of OAciD-MS/MS spectra for ammonium adduct form of TG remains difficult when various unsaturated FAs are contained in the TG molecules. Because the optimal collision energy conditions for obtaining MSL annotations differ widely among lipid subclasses, the collision energy conditions should be changed for profiling the targeted lipid subclasses. Nevertheless, the MS-DIAL 5 program automatically annotates OAciD-MS/MS spectral data, in which an appropriate lipid description is generated based on the quality of the product ion spectrum.

### C=C position-resolved in-depth lipidomics of marmoset brain

Marmosets have attracted considerable attention in neuroscience research because of their neurological and genetic similarities to humans. Untargeted lipidomics has attracted attention for understanding the complexity of brain lipids and their changes during aging ^22–24^; however, little work has been done on the lipid profiling of mammalian brain regions. We applied a structural lipidomics approach to marmoset brain sections using reversed-phase LC (RPLC) coupled to OAciD-MS/MS system, including the frontal lobe, hippocampus, midbrain, cerebellum, and medulla (Tables S3–S5). The total ion chromatograms for each marmoset brain region are shown in Figure S6a. In total, 388 lipids were detected in the marmoset brain, of which 124 molecules were annotated with double bond positions (Figure 3a). The fragmentation behaviors of OAciD-MS/MS, that is, the balance of FA compositions and their C=C positions, were also demonstrated in biological samples, in which most of the PC, PS, and PI molecules were annotated at the structural depth of the double bond positions (Figure 3b). In addition, the double bond positions of sulfatides (SHexCers) on the sphingosine backbone were annotated based on the fragmentation rules of Cers optimized using a standard mixture of EquiSPLASH. The versatility of this fragmentation method was demonstrated by annotating 20 lipid subclasses, including minor lipids such as sphingoid base (SPB), acylhexosylceramides (AHexCers), acylcarnitine (CAR), and monogalactosyl DGs (MGDGs) (Figure 3b and Tables S4 and S5).

**Figure 3.**
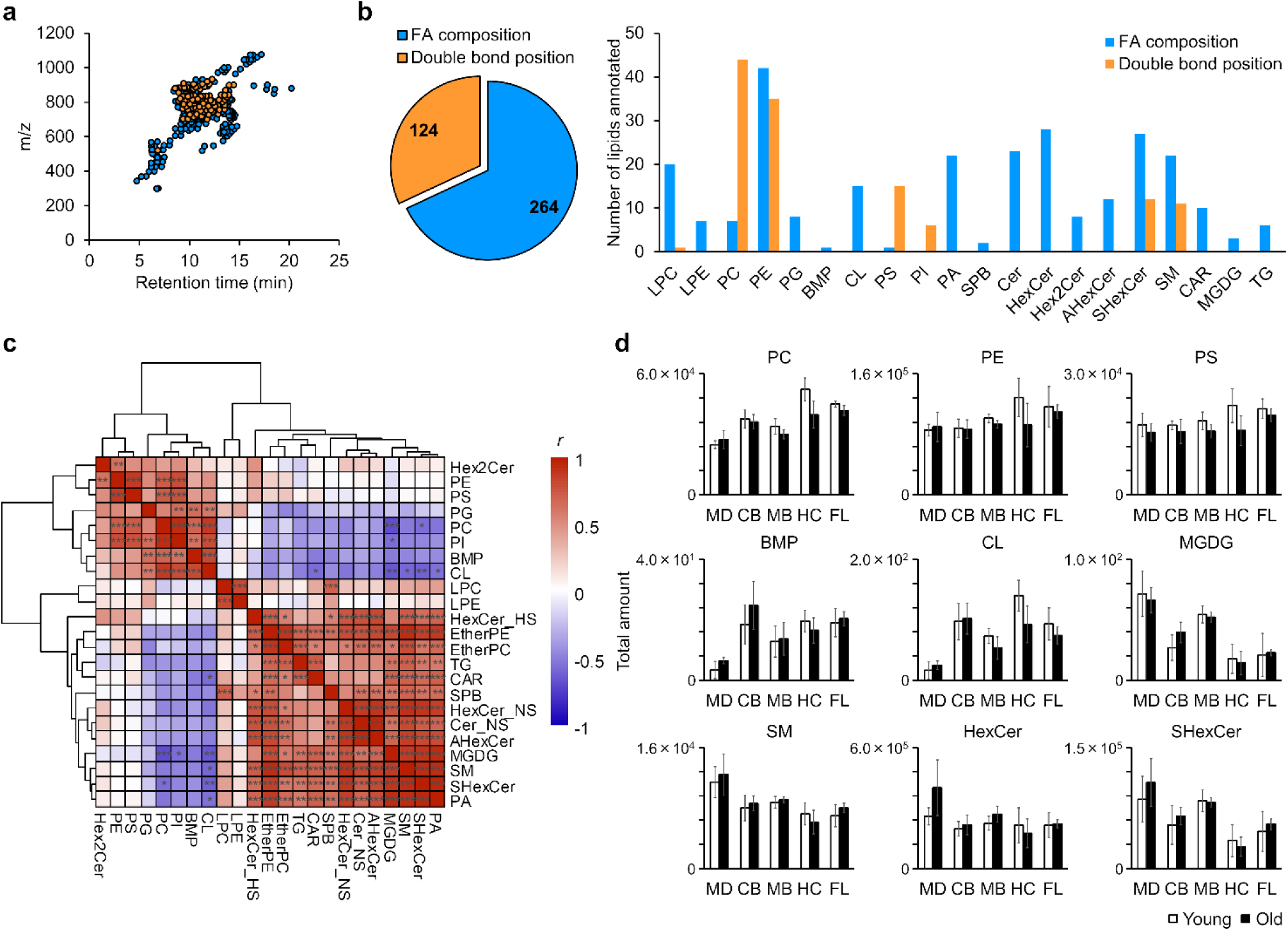
Non-targeted lipidomics of marmoset brain sections. (**a**) Scatter plot of lipid molecules detected in marmoset brain section. (**b**) Number of lipids detected at the structural depth of fatty acyl compositions or double bond positions. (**c**) Hierarchical clustering analysis using the correlation coefficient between lipid subclasses. A total of 40 samples were used for the correlation analysis without discriminating between different regions and ages to characterize lipid subclasses. Correlation analyses were performed using Pearson correlation coefficient. The *p*-values obtained from these analyses were adjusted for multiple comparisons using the false discovery rate method. Adjusted *p*-values were classified into significance levels and displayed using symbols: *** for *p* < 0.001, ** for *p* < 0.01, and * for *p* < 0.05. (**d**) Total amounts of representative lipid subclasses in each region and age. Five regions including frontal lobe (FL), hippocampus (HC), midbrain (MB), cerebellum (CB), and medulla (MD) were analyzed. Error bars indicate the standard deviations of the four biological replicates in each group.

The localization of lipid subclasses in the marmoset brain is similar to that in the mouse brain ^22,23^. Gene expression in the marmoset brain involved in the lipid subclasses was searched using the Marmoset Gene Atlas ^25,26^ (Figure S6b). Hierarchical clustering analysis using the correlation coefficients between lipid subclasses identified two groups consisting mainly of phospholipids and sphingolipids (Figure 3c). Hexosylceramides (HexCers) and SHexCers are enriched in oligodendrocytes that form myelin sheaths on the axons of neurons ^22,23^. These lipid subclasses were slightly higher in the medulla than in other regions (Figure 3d). In addition to these sphingolipids, AHexCers and MGDGs clustered in the same group. MGDGs and galactosylceramides are synthesized from DGs and Cers through catalysis by UDP-galactose-ceramide galactosyltransferase (CGT) ^27^. CGT is expressed in oligodendrocytes and Schwann cells, which is consistent with the high localization of galactosylceramides. *In situ* hybridization of the myelin basic protein (MBP) gene, one of the major components of oligodendrocytes, showed high expression in the midbrain and medulla and low expression in the hippocampus and cerebellum (Figure S6b). The localization of glycolipids, such as SHexCer and MGDG, was similar to that of *Mbp*. Phospholipids, except lysophospholipids, ether PC, ether PE, and phosphatidic acid (PA), clustered in the same group (Figure 3c). Among them, major phospholipids, such as PC, PE, PS, and PI, were correlated, but their specific localization was not observed, suggesting that the phospholipid components of cell membranes were almost constant despite the presence of different cell types. In contrast, low levels of bis(monoacylglycero)phosphate (BMP) and cardiolipin (CL) were observed in the medulla (Figure 3d). BMP, which is synthesized from a lyso-PG (LPG) substrate by ceroid-lipofuscinosis neuronal 5 (CLN-5), is localized in organelles such as late endosomes and lysosomes.^28^ Consistent with this localization, *Cd63*, a late endosome marker, was expressed at low levels in the medulla, as determined by *in situ* hybridization of the marmoset brain (Figure S6b). CL is a specific lipid subclass in the inner mitochondrial membrane that plays structural and functional roles such as maintaining mitochondrial morphology and regulating apoptosis.^29^ A correlation between CL and the mitochondrial integral membrane protein TOMM20 was observed in the rat brain.^29^ High expression of *Tomm20* in the cornu ammonis and dentate gyrus of the hippocampus was observed by *in situ* hybridization (Figure S6b), and high levels of total CLs were detected in the hippocampal region using our lipidomics data (Figure 3d). Consequently, our results indicated that OAciD-MS/MS represented reasonable localization of lipid subclasses reported in previous brain lipidomics study of mouse ^22,23^ and human ^24^.

Several structural isomers, including those at different double bond positions, have been observed in the marmoset brain. Although a single MS1 peak of PC 16:0_16:1 was observed, MS/MS fragments of the C=C position suggested the co-elution of PC 16:0_16:1(Δ7) and PC 16:0_16:1(Δ9) (Figure 4a). Such a co-elution situation is advantageous when using OAciD-MS/MS because the two structural isomers of PC 16:0_16:1(Δ7) and PC 16:0_16:1(Δ9) cannot be elucidated based on EAD-MS/MS (Figure S6c). On the other hand, some MS1 peaks of PC 18:0_22:5 were detected, of which these MS/MS spectra represented PC 18:0_22:5(Δ4,7,10,13,16) and PC 18:0_22:5(Δ7,10,13,16,19), respectively (Figure 4a). Hierarchical clustering analysis of the constituent PC molecules clearly classified structure-dependent groups, such as chain length, odd or even chains, saturated or unsaturated bonds, and double bond positions (Figure 4b). For *n*-9 monounsaturated FAs (MUFAs), the trend differed for chain lengths greater than or less than 20 carbon atoms. PC containing long MUFAs such as PC 18:1(Δ9)_24:1(Δ15), were localized in the medulla, cerebellum, and midbrain (Figure S6d). Among the *n*-6 PUFAs, PCs containing arachidonic acid (20:4, *n*-6), docosatetraenoic acid (22:4, *n*-6), and docosapentaenoic acid (22:5, *n*-6) were clustered in the same group; however, they did not correlate with those containing linoleic acid (18:2, *n*-6), even though the constituent FAs were placed in the same FA metabolic pathway. Different patterns of fatty acyl composition and specific localization of some constituent molecules have been determined in the marmoset brain. Although eicosapentaenoic acid (EPA) (20:5, *n*-3) and docosahexaenoic acid (DHA) (22:6, *n*-3) are in the same synthetic pathway as *n*-3 PUFAs, EPAs levels are very low compared to DHAs in the brain. Only PC 16:0_20:5(Δ5,8,11,14,17) was detected as an EPA-containing phospholipid also in the marmoset brain (Table S5). The localization of phospholipids was mainly dependent on the fatty acyl compositions rather than on the lipid subclasses. High levels of saturated PC and PE were detected in the frontal lobe, and low levels were detected in the medulla (Figure S6d). The most specific localization was observed for phospholipids containing diacyl PUFAs. Phospholipids containing diacyl *n*-3 PUFAs are localized in the cerebellum where the amounts of PS 22:5(Δ7,10,13,16,19)_22:6(Δ4,7,10,13,16,19) were especially enriched when compared with the other tissues (Figures 4c and S6e).

**Figure 4.**
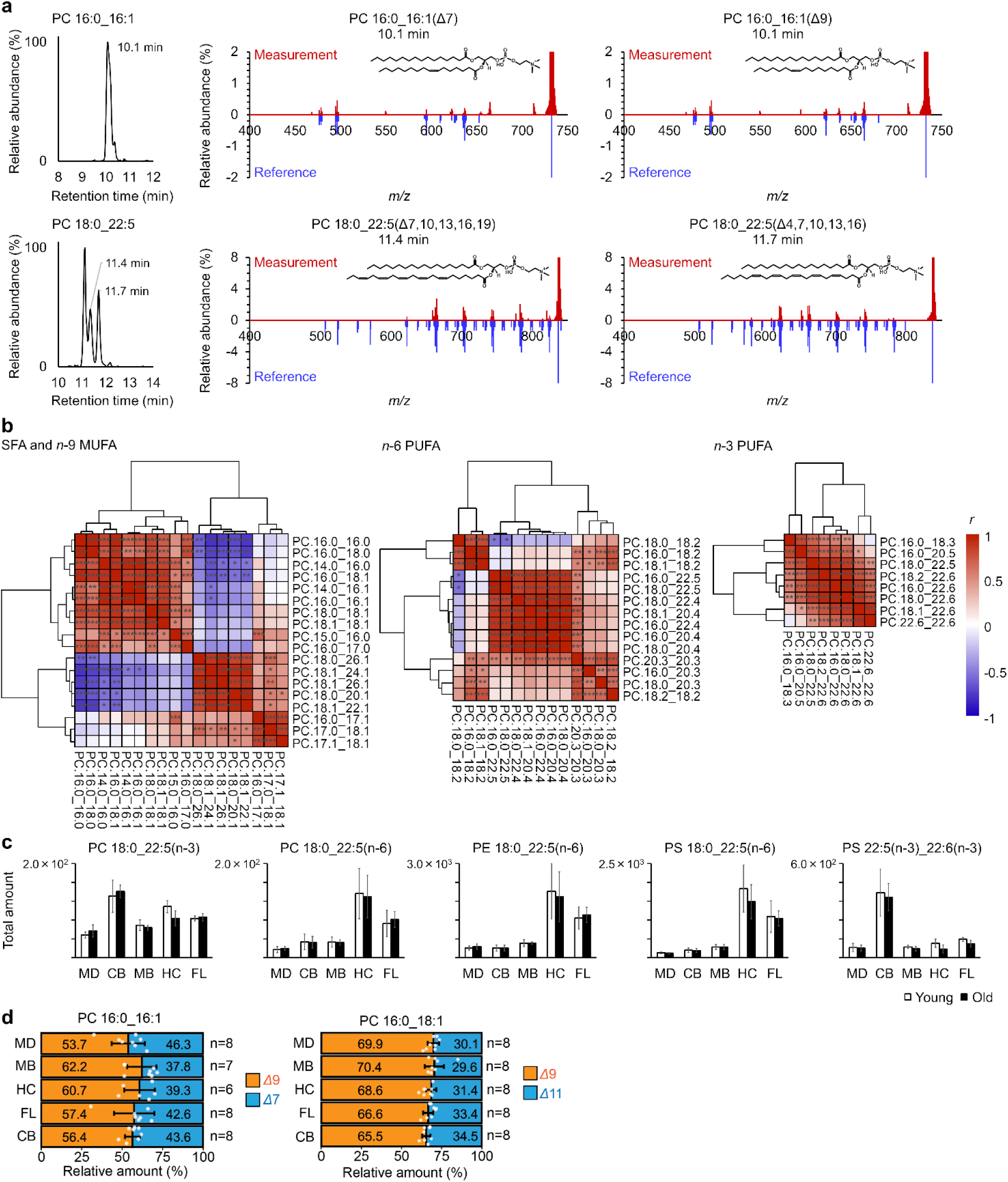
Characterization of lipid molecules at the structural depth of the double bond position. (**a**) Chromatograms and MS/MS spectra of structural isomers with different C=C positions. (**b**) Hierarchical clustering analysis using the correlation coefficient between PC molecules. Saturated FA (SFA) and *n*-9 MUFA, *n*-6 PUFA, and *n*-3 PUFA were analyzed using the same methods as in Figure 3c. (**c**) Total amounts of constituent molecules in each region and age. Error bars indicate the standard deviations of the four biological replicates in each group. (**d**) The ratios of PC 16:0_16:1(Δ7) and PC 16:0_16:1(Δ9) and ratio of PC 16:0_18:1(Δ9) and PC 16:0_18:1(Δ11) among tissues. Error bars indicate the standard deviations of the biological replicates in each group where both young– and aged marmosets were included. Due to the limit of detection in the data-dependent acquisition mode for PC 16:0_16:1, total three samples containing one midbrain and two medulla samples were excluded.

Finally, we characterized the ratios of PC 16:0_16:1(Δ7) to PC 16:0_16:1(Δ9) and PC 16:0_18:1(Δ9) to PC 16:0_18:1(Δ11) using data-dependent OAciD-MS/MS spectral data (Figure 4d). According to our previous study,^20^ the C=C position-specific fragmentation efficiency between PC 16:0_18:1(Δ9) and PC 16:0_18:1(Δ11) was nearly equal, allowing the relative abundance of these lipid isomers to be estimated by MS/MS peak heights. The product ion peak heights of *m/z* 636.42 and *m/z* 664.45 were used to calculate the ratio of PC 16:0_16:1(Δ7) to PC 16:0_16:1(Δ9), while the peaks of *m/z* 664.45 and *m/z* 692.49 were used for the ratio of PC 16:0_18:1(Δ9) to PC 16:0_18:1(Δ11). As a result, the ratios of these isomers were approximately equal across the marmoset tissues examined, with PC 16:0_16:1(Δ9) and PC 16:0_18:1(Δ9) identified as the major isomers. FA 16:1(Δ9) and FA 18:1(Δ9) are biosynthesized by stearoyl coenzyme A desaturase 1 (SCD1) using palmitic acid (16:0) and stearic acid (18:0) as substrates, respectively. FA 18:1(Δ11), known as cis-vaccenic acid, is biosynthesized by ELOVL fatty acid elongase 6 (ELOVL6) using 16:1(Δ9) as a substrate. The FA 16:1(Δ7) isomer, rarely reported in mammalian cell studies, may be synthesized through the beta-oxidation of FA 18:1(Δ9). Given that the brain is a mitochondria-rich organ, beta-oxidation activity is likely higher compared to other organs, though the mechanistic role of PC 16:0_16:1(Δ7) in the brain remains unknown.

## Discussion

In this study, we developed an advanced structural lipidomics method using OAciD-MS/MS to determine the C=C position in the fatty acyl chains of diverse lipid subclasses. Our OAciD-MS/MS-based approach overcame technical limitations related to lipid subclass coverage and analytical throughput by enabling analysis in negative ion mode and dual fragmentation of CID and OAD. Notably, the method eliminates the need for data acquisition of conventional CID lipidomics by achieving simultaneous cleavage of CID and OAD through residual H_2_O vapor and radical species (O and/or OH·). The OAD-MS/MS system offers two key advantages over previous methods for determining double bond positions. First, the OAD reaction is independent of ion mode polarity (positive or negative), though sensitivity varies depending on lipid subclasses and adduct ions. OAciD-MS/MS can analyze a wide range of lipids with diverse chemical properties, reducing the reliance on sodium adduct ions, which can contaminate mass spectrometers. Second, the C=C position is easily annotated, even when isomers co-elute at the MS1 level, because the product ions related to the double bond are limited. For example, when using EAD-MS/MS, the entire MS/MS spectrum must be fingerprinted for lipid annotation (Figure S6c). This allows quantification of individual lipid molecules using product ions, even when isomers are not baseline-separated by LC.

Although the OAD-MS/MS fragmentation mechanism has been demonstrated for lipidomics, reports on its application to hydrophilic metabolites remain sparse. Unlike conventional MS/MS, the OAD-MS/MS system detects O– and O_2_-attached precursor ions, with fragment ions formed by oxidation at the C=C position. While microwave discharge of ultrapure H_2_O produces hydrogen radicals (H·), no H·-specific fragment ions were observed from the cleavage of carbon-carbon single bonds.^13^ Consistent with previous reports, H· did not contribute to covalent bond dissociation. Instead, fragment ions of fatty acyl chains at the C=C position are induced by oxidation with O and OH·. For instance, neutral losses of 96.13 Da (between Δ10 and Δ11 positions) and 138.14 Da (between Δ8 and Δ9 positions) were observed for the FA 18:1(Δ9) chain. The 96.13 Da loss occurs when OH· adds to the Δ10 position, forming a double bond at Δ9-Δ10 through electron movement (Figure 1a).^18^ Conversely, the 138.14 Da loss is initiated by the addition of O at the Δ9 position, leading to cleavage between the Δ8 and Δ9 positions. The resulting fragment ion then adds H· at the radical site (Figure 1a).^18^ These major fragments were detected independently of lipid subclass and MS polarity.

Several brain morphologies, including cortical thinning, Purkinje and granule cell loss in the anterior cerebellar lobe, neuronal loss and/or decreased neuronal density in the hippocampus, are altered by aging.^30–32^ Recently, marmoset brain databases, such as two– and three-dimensional brain atlases,^33^ the prefrontal cortex connectome,^34^ and *in situ* hybridization gene atlases,^25,26^ have been developed. However, research on the metabolome of mammalian brains is still in its early stages.^24^ Brain lipidomics, with its structural depth and spatiotemporal localization, offers potential for characterizing altered lipid metabolism in diseases.^35^ Previous studies using untargeted lipidomics in mice have revealed relationships between specific lipid metabolisms and aging, such as BMP and sulfonolipids in various organs and glycolipids in the kidney.^36^ Although the lipid metabolites related to aging were not identified in this study due to the small sample size, lipid localization was assessed at a deep structural level, including double bond positions. Hierarchical clustering analysis revealed that localization patterns were primarily dependent on fatty acyl chain structures, including chain length, number of double bonds, and positions. Lipid metabolism is organized into layers of *de novo* synthesis of lipid subclasses (Kennedy pathway) and remodeling of fatty acyl chains (Lands cycle). A previous study suggested that *n*-3 very long-chain PUFA and DHA (22:6, *n*-3) are not incorporated into phospholipids via the same metabolic pathways.^20^ Thus, the complete mechanisms of *de novo* synthesis and remodeling pathways in phospholipids remain unclear. While we are just beginning to investigate the localization patterns of individual lipid molecules in the brain, structural lipidomics will provide new insights into complex lipid metabolism. In addition to its structural depth, the untargeted lipidomics method and MS-DIAL 5 software characterized minor lipid subclasses, including AHexCer and MGDG, despite applying dual CID and OAD modes with H_2_O vapor. This study demonstrated the potential of OAciD-MS/MS for next-generation lipidomics by analyzing double bond positions while detecting minor lipids in the marmoset brain.

Despite the enhanced structural depth achieved through this novel fragmentation method, challenges remain in the annotation and quantification of constituent molecules. The fragmentation efficiency at double bond positions is less than 5% of the precursor ion, making the dynamic range of these fragment ions too narrow for effective annotation. Further development in hardware, such as data acquisition modes, and software, such as MS-DIAL, is needed to integrate data with both low and high collision energy settings, allowing simultaneous detection of fragments related to fatty acyl composition and double bond position. This would improve deep annotation, especially for Cer and DG, whose C=C positions are difficult to annotate at high energies. From a pretreatment perspective, enriching specific lipid subclasses is necessary to maximize the dynamic range. We have recently developed a solid-phase enrichment protocol to enhance the dynamic range, resulting in the characterization of minor lipid subclasses, such as very long-chain PUFA-containing Cer, HexCers, and dihexosylceramides (Hex2Cers) in testis, mono– and dihexosyl monoacylglycerols in feces, and acetylated and glycolylated derivatives of gangliosides using the conventional CID-MS/MS system.^37^ This protocol has contributed to improved lipid annotations. Furthermore, low-flow LC (<10 μL/min) can accelerate high-sensitivity lipidomics, although the sensitivity of C=C position-related fragment ions in dual-MS/MS spectra in negative ion mode remains a challenge. Despite these technical hurdles, this next-generation lipidomics approach could shed light on complex lipid metabolism, including the remodeling of fatty acyl chains.

## Methods

### Chemicals and Reagents

Reagent grade chloroform (CHCl_3_), 1 mol/L ammonium acetate solution for high-performance LC, acetonitrile (MeCN), methanol (MeOH), 2-propanol (IPA), and ultrapure H_2_O for quadrupole time-of-flight MS (QTOFMS) were obtained from FUJIFILM Wako Pure Chemical Corp. (Osaka, Japan). Individual lipid synthetic standards, EquiSPLASH containing equal concentrations of deuterium-labeled lipid standard (each, 100 μg/mL), and UltimateSPLASH containing 69 deuterium-labeled synthetic lipids at various concentrations were purchased from Avanti Polar Lipids Inc. (Alabaster, AL, U.S.A.).

### Animals

The experimental protocols for the animal experiments were approved by RIKEN’s Wako Animal Experiments Committee. Four young (1.5–1.9 years old, 3 males and 1 female) and four old (14.4–16.3 years old, all males) common marmosets were purchased from the RIKEN Research Resources Division and CLEA Japan, Inc. (Tokyo, Japan), respectively (Table S3). After euthanasia with an overdose of ketamine (50 mg/kg, i.m.), the medulla, cerebellum, and midbrain were sequentially removed from the dissected brain, and the remaining brain was divided at the midline. Hippocampus and frontal lobe were collected from the right brain. Brain tissues were harvested and immediately frozen after dissection and stored at −80°C until lipid extraction.

### Liquid-liquid extraction

The Bligh and Dyer method was used as previous method.^38^ Marmoset brain sections were lyophilized overnight in the dark. After the brain tissue was crushed using a ball mill, an average volume of 5 mg was transferred to a clean tube. Samples were mixed with 1,000 μL of MeOH/CHCl_3_/H_2_O (10:4:4, v/v/v). Lipids were extracted using a vortex mixer for 1 min and then ultrasonicated for 5 min. The solution was centrifuged at 16,000 g for 5 min at 4°C, and 700 μL of the supernatant was transferred to a clean tube. The supernatant was mixed with 235 μL of CHCl_3_ and 155 μL of H_2_O using a vortex mixer for 1 min. Both layers were collected after centrifugation at 16,000 g for 15 min at 4°C. Lipid extracts were dried using a centrifuge evaporator. They were dissolved with MeOH containing 200 times diluted EquiSPLASH mixture (each, 500 ng/mL).

### High-throughput analysis for method optimization

The LC/QTOFMS system was composed of a Nexera and an LCMS-9030 equipped with an OAD RADICAL SOURCE I (Shimadzu Corp., Kyoto, Japan). A high-throughput analytical system was used to efficiently optimize the method without causing an MS shift or change in sensitivity owing to the long period of operation.^39^ The LC conditions were as follows: injection volume, 1 μL; mobile phase, MeCN/MeOH/H_2_O (1:1:3, v/v/v) (A) and MeCN/IPA (1:9, v/v) (B) (both contained 10 nM of ethylenediaminetetraacetic acid (EDTA) and 5 mM of ammonium acetate); flow rate, 600 μL/min; column; Unison UK-C18 MF (50 × 2.0 mm, 3 μm, Imtakt Corp., Kyoto, Japan); gradient, 0.1% (B) (0.1 min), 0.1–15% (B) (0.1 min), 15–30% (B) (0.9 min), 30–48% (B) (0.3 min), 48–82% (B) (4.2 min), 82–99.9% (B) (1.3 min), 99.9% (B) (0.2 min), 99.9–0.1% (B) (0.1 min), 0.1% (B) (1.4 min); column oven temperature, 65°C. The column eluent was mixed with 10–50 μL/min of 50% methanol containing 200 μM sodium acetate post-column when forming the sodium adduct ion (Figure S1a). MS conditions were as follows: nebulizing gas flow, 3.0 L/min; heating gas flow, 10.0 L/min; drying gas flow, 10.0 L/min; interface temperature, 300°C; DL temperature, 250°C; heat block temperature, 400°C; CID gas, 17 kPa (using H_2_O and its radicals) or 230 kPa (using Ar gas); TOF range; *m/z* 75–1250; event time; 200 ms (MS scan) or 100 ms (MS/MS); interface voltage, 4 kV (positive ion mode) or –4 kV (negative ion mode); Q1 resolution, 4 Da. The precursor ions of the product ion scan used to investigate the fragmentation efficiency between positive and negative ion modes and to optimize the collision energy are summarized in Table S1.

### Lipidome analysis of the marmoset brains

The LC conditions were as follows: injection volume, 2 μL; mobile phase, MeCN/MeOH/H_2_O (1:1:3, v/v/v) (A) and MeCN/IPA (1:9, v/v) (B) (both contained 10 nM of EDTA and 5 mM of ammonium acetate); flow rate, 300 μL/min; column; Unison UK-C18 MF (50 × 2.0 mm, 3 μm, Imtakt Corp., Kyoto, Japan); gradient, 0.5% (B) (1 min), 0.5–40% (B) (4 min), 40–64% (B) (2.5 min), 64–71% (B) (4.5 min), 71–82.5% (B) (0.5 min), 82.5–85% (B) (6.5 min), 85–99% (B) (1 min), 99% (B) (2 min), 99–0.5% (B) (0.1 min), 0.5% (B) (2.9 min); column oven temperature, 45°C. LCMS-9030 conditions were the same as those used for method optimization.

Structural analysis of PC 16:0_16:1 and PC 16:0_18:1 was performed using a ZenoTOF 7600 (SCIEX, Framingham, MA, USA). ZenoTOF 7600 conditions were used as previous method.^20^ MS parameters were as follows: ion source gas 1, 40 psi; ion source gas 2, 80 psi; curtain gas, 30 psi; CAD gas, 7; temperature, 250°C; polarity, positive ion mode; spray voltage, 5500 V; declustering potential, 80 V; collision energy, 10 V; scan range; *m/z* 75–1250 (MS1) and 100–1250 (MS/MS); accumulation time, 200 ms (MS1) and 100 ms (MS/MS); time bins to sum, 4 (MS1) and 8 (MS/MS); electron beam current, 7000 nA; zeno threshold, 1000000 cps; kinetic energy, 14 and 18 eV; EAD RF, 150 Da; reaction time, 98 ms.

### Data analysis

In the method development, peak area was calculated using LabSolutions Insight LCMS (Shimadzu). For the lipidome analysis of marmoset brains, lipid molecules were annotated using MS-DIAL version 5.1. Representative parameters were as follows: minimum peak height for peak picking, 100; retention time tolerance for peak alignment, 0.15 min; MS1 tolerance, 0.01 Da; MS2 tolerance, 0.025 Da. The other data processing parameters and adduct ions used for each lipid subclass are listed in Table S6 and S4, respectively. The annotated results were manually curated using the retention times and MS/MS spectra of the peaks in both positive and negative ion modes. The data matrix was normalized based on the dry weight of the marmoset brain tissue and the ion abundance of the EquiSPLASH mixture. The lipidome results are summarized in Table S5.

### Lipid nomenclature and quantification

The nomenclature used in this manuscript is described in accordance with the Lipidomics Standards Initiative.^40^ For example, PC 16:0_18:1 indicates that the individual fatty acyl chains was determined by MS/MS. The detailed position is described as PC 16:0_18:1(Δ9) when fragment ions derived from the C=C positions are obtained by OAciD. In contrast, the type of quantification corresponded to level 3 (lipid subclass other than analyte or no co-ionization of analyte and internal standard) of the Lipidomics Standards Initiative International Guidelines. The corresponding internal standards used to quantify each lipid subclass are summarized in Table S4. The quantitative performance characteristics of the RPLC/MS system were discussed in a previous report^37^.

### Data availability

All raw MS data are available on the RIKEN DROPMet website (https://prime.psc.riken.jp/menta.cgi/prime/drop_index) under the index number DM0066. The MS-DIAL source code is available at https://github.com/systemsomicslab/MsdialWorkbench.

## Acknowledgments

We thank the staff of the Research Resources Division, RIKEN Center for Brain Science, for the management and breeding of some of the marmosets. This research was supported by the Japan Agency for Medical Research and Development (AMED) under Brain Mapping by Integrated Neurotechnologies for Disease Studies (Brain/MINDS) (JP15dm0207001 to H.K. and H. Tsugawa), the Japan Science and Technology Agency (JST) Exploratory Research for Advanced Technology (ERATO) (JPMJER2101 to H. Tsugawa), the JSPS KAKENHI (24K02011, 24H00043, 24H00392, 24K21269 to H. Tsugawa), JST National Bioscience Database Center (JPMJND2305 to H. Tsugawa), and Technologically Advanced research through Marriage of Agriculture and engineering as Groundbreaking Organization (TAMAGO to H. Tsugawa).

## Author Contributions

H. Takeda, H.K. and H. Tsugawa designed the research. M.O. and H. Takahashi developed the hardware of OAD-MS/MS system. B.B. and H. Tsugawa developed the MS-DIAL software. H. Takeda and M.O. optimized the method, analyzed lipids, and performed peak annotation and quantification. H.K. H.O. and H.K. prepared the animals and N.K. dissected them. H. Takeda and H. Tsugawa wrote the manuscript. H. Takeda, B.B. and H. Tsugawa prepared the figures. All authors have given approval to the final version of the manuscript.

