## Supporting information for "Dual fragmentation via collision-induced and oxygen attachment dissociations using water and its radicals for C=C position-resolved lipidomics"

#### Corresponding Authors

#### Contents

Supplementary Figure 1–6

Supplementary Table 1–6

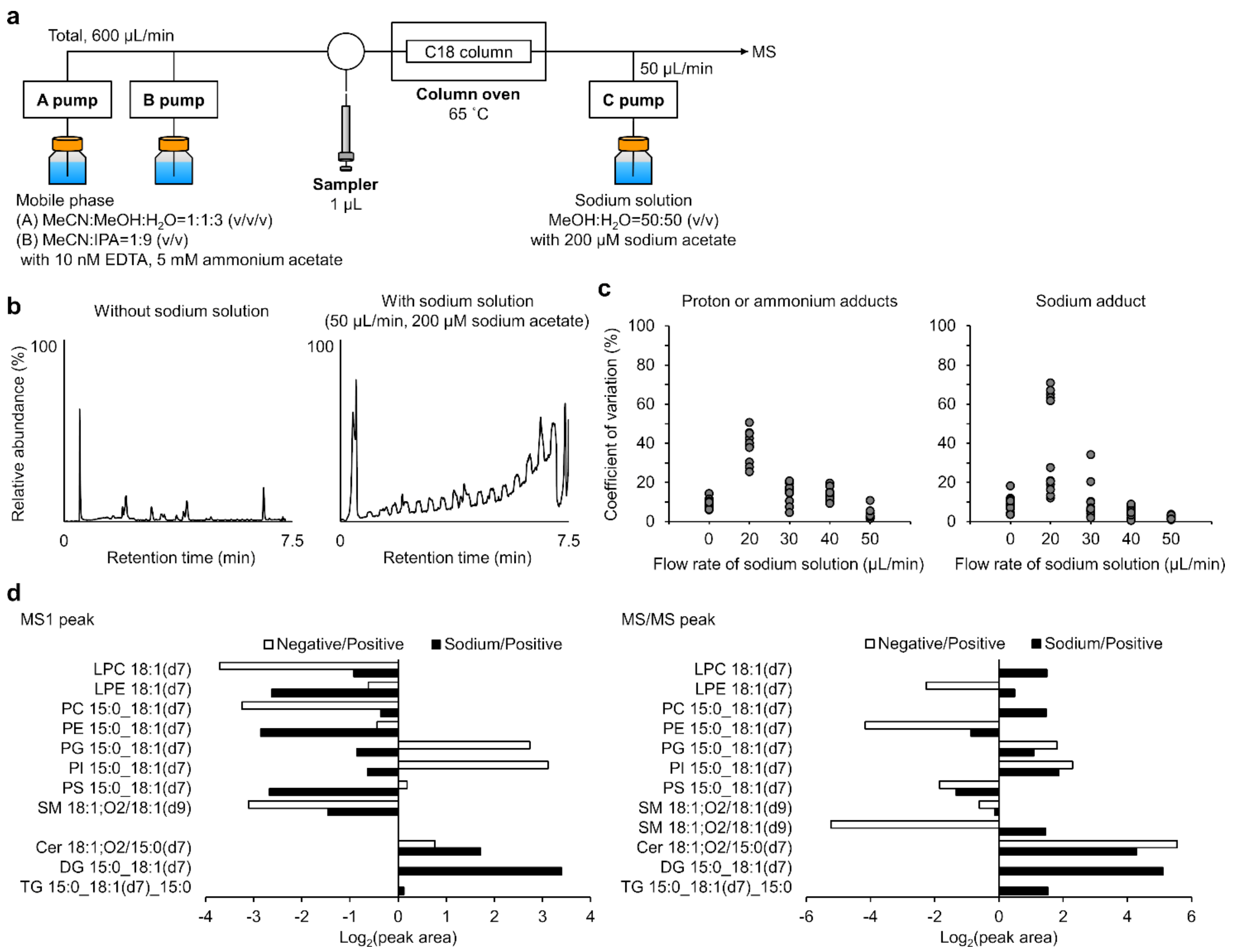


**Figure S1. Structural analysis of lipid synthetic standards using sodium adduct.** (**a**) Schematic of the LC system for adding the sodium solution. (**b**) Total ion chromatogram of LC systems without or with sodium solution. (**c**) Coefficient of variation at each flow rate of sodium solvent. Each plot shows the synthetic standards in the EquiSPLASH mixture. For all proton, ammonium, and sodium adduct ions, 50 µL/min of sodium solvent represents high reproducibility. (**d**) Comparison of the sensitivity for each adduct ion by calculating the MS1 and MS2 peak areas. The positive and negative were analyzed using the conventional LC system without sodium solution. The sodium adduct was formed by confluence of the column eluent and sodium solution. The MS2 peaks were obtained using the product ion scan mode (Table S1).

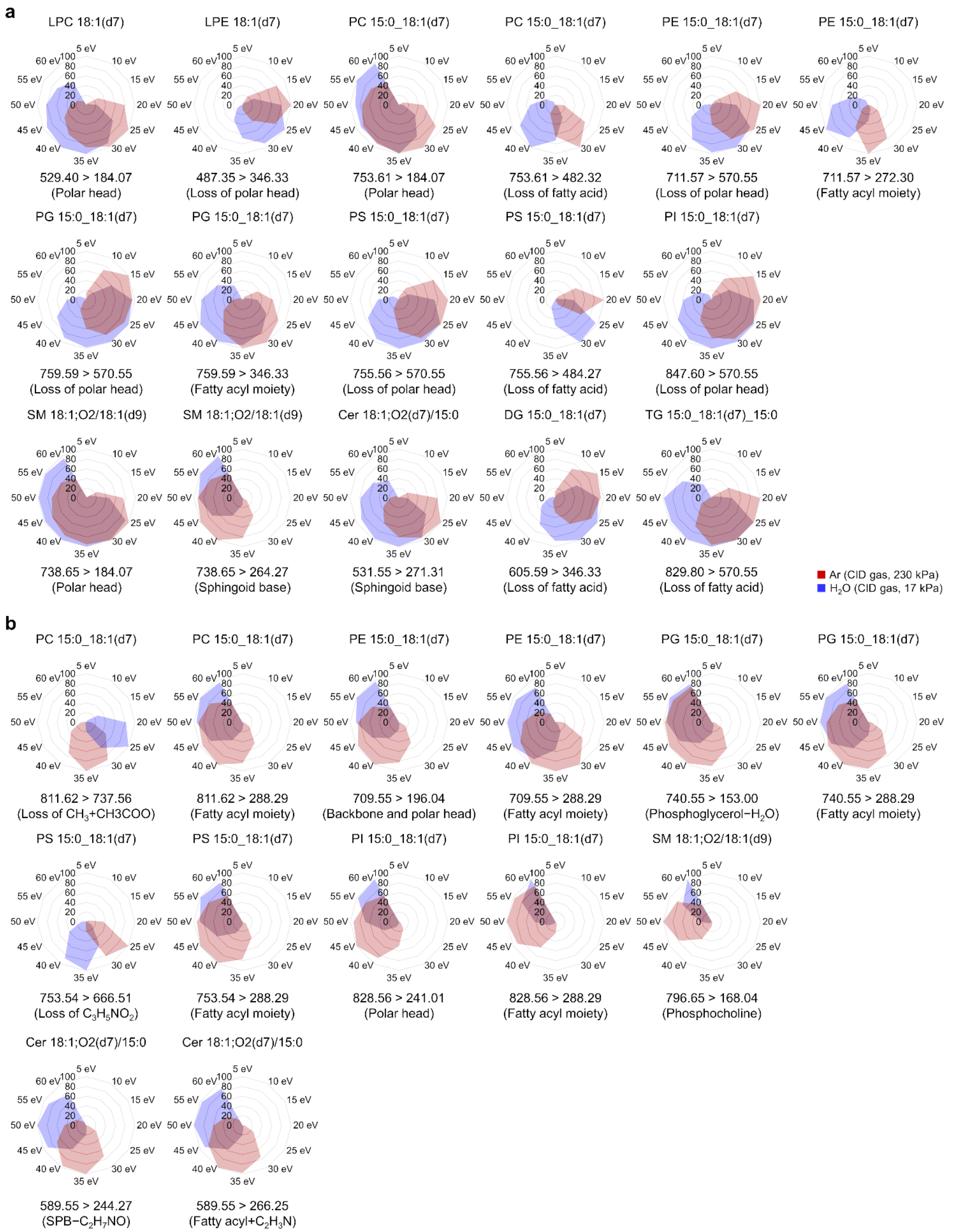


**Figure S2. Comparison of fragmentations between Ar gas and $H_2O$ vapor.** The area value of MS2 peak was normalized to the maximum peak detected under the collision energy range of 5–60 eV. (**a**) Results in positive ion mode. (**b**) Results in negative ion mode.

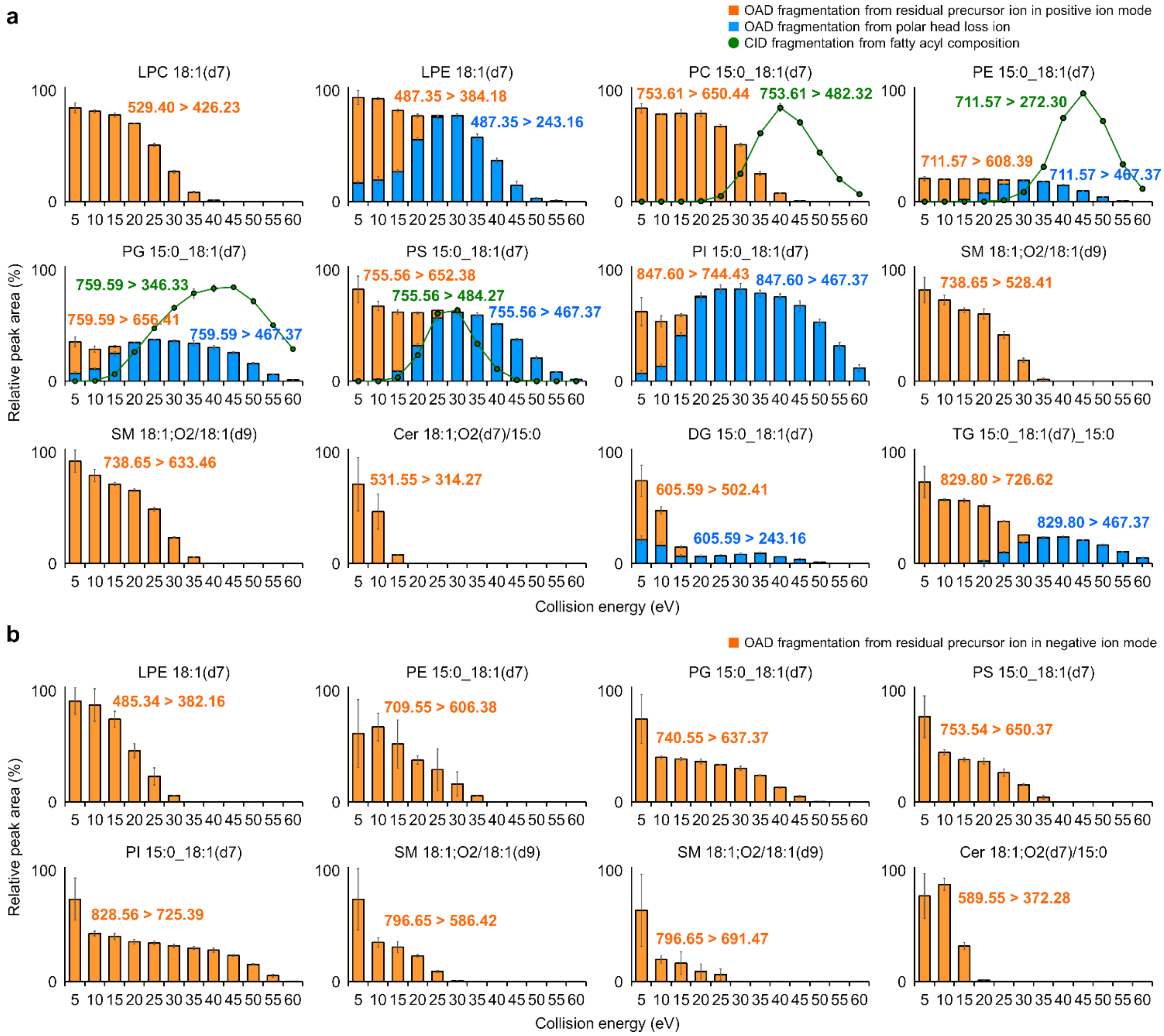


**Figure S3. Fragmentation at the C=C position by increasing the collision energy.** The orange and blue bars represent the MS2 peak area of oxidative fragments at the C=C position of residual precursor and polar head loss ions, respectively. The green line plot shows the fragments at the fatty acyl chains, also observed in CID mode using Ar gas. Error bars indicate the standard deviations of the four analytical replicates. (**a**) Results in positive ion mode. (**b**) Results in negative ion mode.

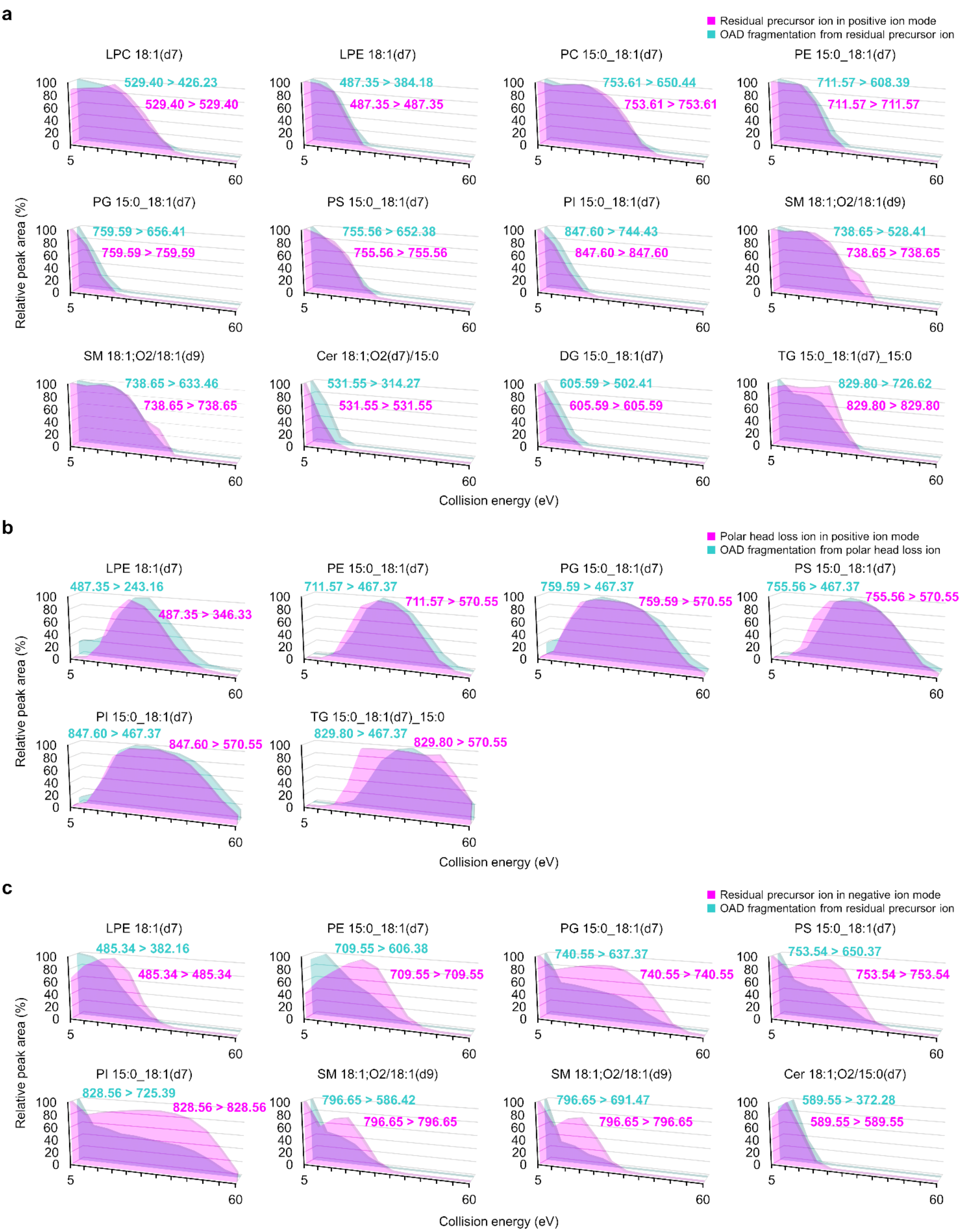


**Figure S4. Fragmentation at the C=C position by increasing the collision energy.** The MS2 peak area of fragments is shown at the C=C position from residual precursor or polar head loss ions. Reactants and their OAD fragments at the C=C position are shown in pastel pink and blue, respectively. Oxidative fragments at the C=C position of residual precursor ions in positive ion mode (**a**), polar head loss ions in positive ion mode (**b**), and residual precursor ions in negative ion mode (**c**) are summarized.

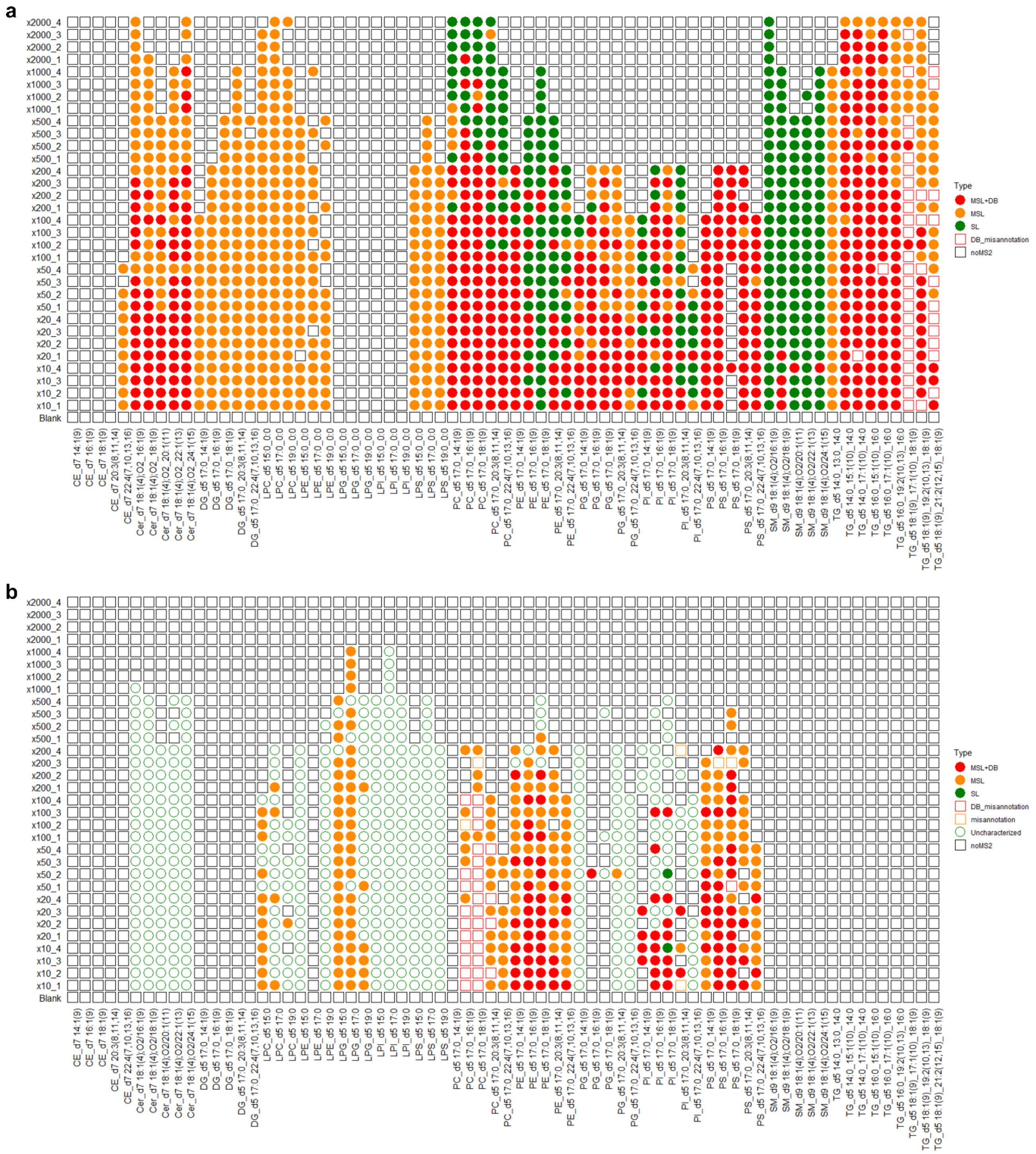


**Figure S5. Annotated results of fatty acyl compositions and their double bond positions using MS-DIAL 5.** When compared to Figure 2, the annotation results of individual samples were described. The definitions of color and symbol are the same as used in Figure 2c. (**a**) Results in positive ion mode. (**b**) Results in negative ion mode.

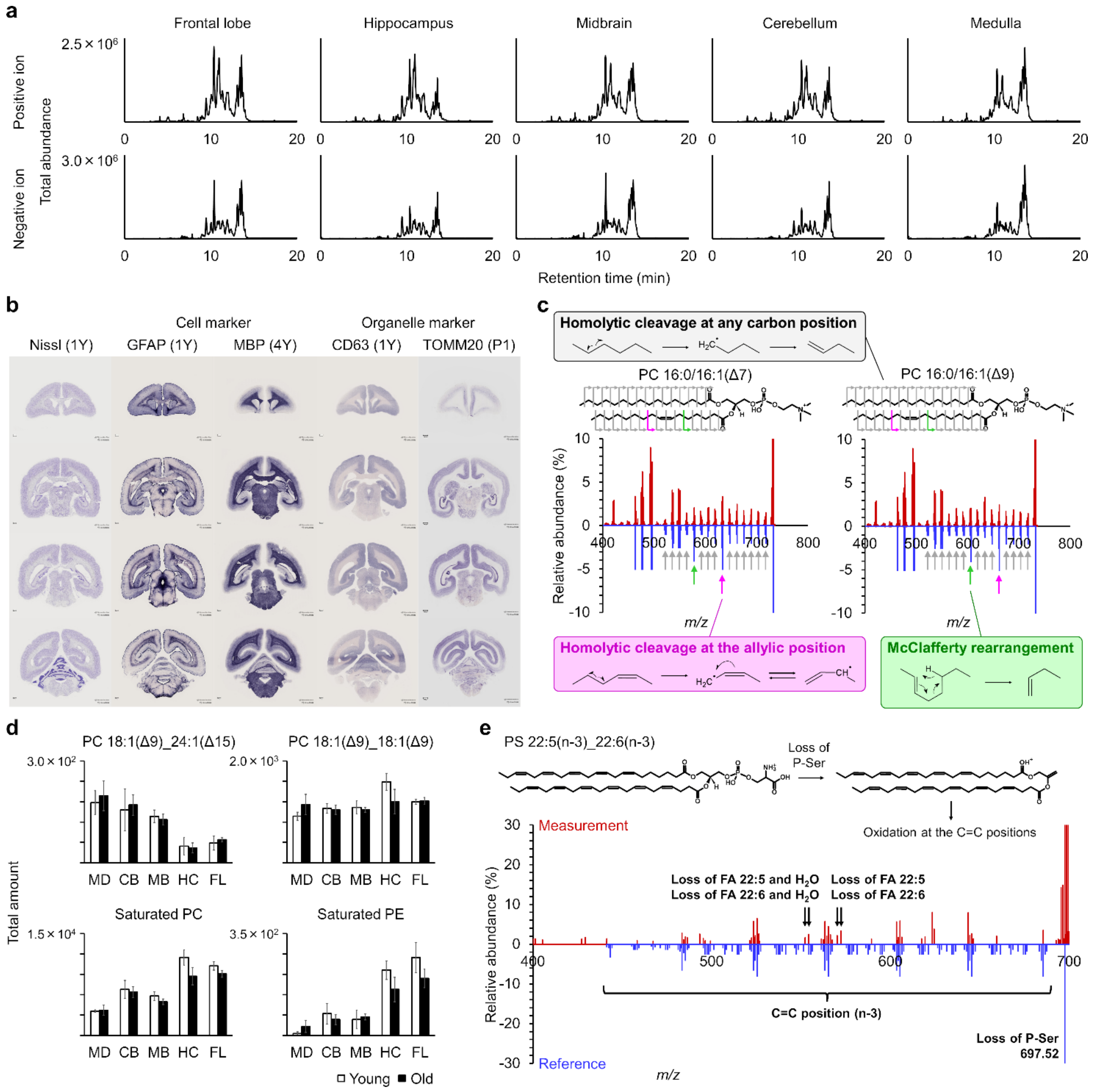


**Figure S6. Double bond-resolved in-depth lipidomics of marmoset brain.** (**a**) Total ion chromatogram of five sections of marmoset brain. (**b**) *In-situ* hybridization of cell and organelle markers in marmoset brain. The images were downloaded from the *Marmoset Gene Atlas* (https://gene-atlas.brainminds.jp/).[25,26] (**c**) Structural analysis of PC 16:0_16:1 using EAD-MS/MS. The peak of PC 16:0_16:1 at 10.1 min in Figure 4a was targeted. The double bond positions can be annotated with an allyl radical product and a hydrogen loss fragment produced by McLafferty rearrangement.[17] The lower blue mass spectra represent the reference library implemented in the MS-DIAL 5 software.[20] (**d**) Total amounts of *n*-9 MUFA-containing PC and saturated PC and PE in each region and age. Error bars indicate the standard deviations of the four biological replicates in each group. (**e**) Structural analysis of PS 22:5_22:6 using OAciD-MS/MS.

### Supplementary Tables

**Table S1. Product ion scan of deuterium labeled EquiSPLASH standards.**

**Table S2. Calibration curve of RPLC/OAciD-MS system using EquiSPLASH standard.**

**Table S3. Sample information of individual marmoset.**

**Table S4. Summary of lipid subclasses and their adduct ions used to calculate lipid amounts in the study.**

**Table S5. Lipidome analysis of marmoset brain.** Lipidomics data were normalized using the sample volume (approximately 5 mg) and ion abundance of the internal standard added after lipid extraction by the modified Bligh and Dyer method (see Tables S2 and S3).

**Table S6. Parameters of MS-DIAL software.**
